# Efficient *in situ* barcode sequencing using padlock probe-based BaristaSeq

**DOI:** 10.1101/180323

**Authors:** Xiaoyin Chen, Yu-Chi Sun, George M Church, Je Hyuk Lee, Anthony M Zador

**Affiliations:** Cold Spring Harbor Laboratory, Cold Spring Harbor, NY 11724, USA; Wyss Institute, Harvard Medical School, Boston, Massachusetts, USA.; Department of Genetics, Harvard Medical School, Boston, Massachusetts, USA

## Abstract

Cellular DNA/RNA tags (barcodes) allow for multiplexed cell lineage tracing and neuronal projection mapping with cellular resolution. Conventional approaches to reading out cellular barcodes trade off spatial resolution with throughput. Bulk sequencing achieves high throughput but sacrifices spatial resolution, whereas manual cell picking has low throughput. *In situ* sequencing could potentially achieve both high spatial resolution and high throughput, but current *in situ* sequencing techniques are inefficient at reading out cellular barcodes. Here we describe BaristaSeq, an optimization of a targeted, padlock probe-based technique for *in situ* barcode sequencing compatible with Illumina sequencing chemistry. BaristaSeq results in a five-fold increase in amplification efficiency, with a sequencing accuracy of at least 97%. BaristaSeq could be used for barcode-assisted lineage tracing, and to map long-range neuronal projections.

**Key Points:**

- *In situ* sequencing by gap-filling padlock probes is limited by the strand displacement of DNA polymerases
- Illumina sequencing chemistry offers superior signal-to-noise ratio *in situ* compared to sequencing by ligation
- BaristaSeq as an accurate method for barcode sequencing *in situ* with improved gap-filling efficiency

## Introduction

DNA/RNA barcodes can be used to uniquely label individual cells within a population for lineage tracing(1–5) or neuronal projection mapping(6). Conventional approaches to reading out cellular barcodes trade off spatial resolution with throughput. Bulk sequencing achieves high throughput but sacrifices spatial resolution, whereas manual cell picking is low throughput. *In situ* hybridization approaches preserve cellular location but even highly multiplexed approaches cannot readily distinguish barcodes with a diversity in excess of 10^2^-10^3^, whereas in some applications barcode diversity can be 10^6^-10^9^ or beyond (7,8).

To overcome these limitations, we have been developing an *in situ* sequencing approach to read out cellular barcodes. *In situ* sequencing (9,10), which provides information at sub-cellular resolution, has the potential to achieve both high spatial resolution and high throughput. Because the diversity of a random barcode library scales exponentially with the length of the barcode, *in situ* sequencing of even 25 nucleotides is sufficient to distinguish 4^25^ = 10^15^ barcodes, which is adequate for most applications.

There are currently three approaches for amplifying and sequencing mRNAs *in situ* with distinct advantages and disadvantages (Fig. 1). Each begins by reverse transcribing the target into cDNA. In the first approach, the no-gap padlock approach (11) (Fig. 1A), a padlock probe is hybridized to the cDNA. A padlock probe is an oligonucleotide with two “arms” complementary to two adjacent sequences on the target, linked by a generic backbone sequence between the two arms. The arm hybridizing downstream, however, is placed at the 5’ end of the padlock probe, allowing the two arms to be ligated using a double-stranded DNA ligase to form a circularized ssDNA. The circularized padlock probe is then used as a template for rolling circle amplification (RCA), producing a rolling circle colony, or “rolony”—a <1 μm nanoball of DNA that consists of thousands of copies of the original sequence (Fig. 1A). In this no-gap padlock approach, the barcode is built into the backbone of the padlock probes to differentiate among padlocks targeting different mRNAs(12). Compared with the other *in situ* sequencing approaches, this approach has the advantage of higher sensitivity with ~30% RNA detection(12), but has the disadvantage that it does not capture the actual sequence of the targeted RNA since only the padlock probe sequence, not the cDNA sequence, is amplified during RCA. Because a unique probe must be synthesized for each targeted gene, this approach is not suitable for sequencing high-diversity barcode libraries. The second approach, the gap padlock approach (9) (Fig. 1B), also relies on padlocks, but captures the actual sequence of the target barcode. In this approach, the padlock probes have a gap between the two arms, and a DNA polymerase is used to fill this gap (i.e. gap-filling) before circularization of the padlock probe. This process copies a part of the cDNA into the RCA template (Fig. 1B). The copied barcode is then sequenced *in situ*. Because the DNA/RNA barcodes are flanked by known sequences, the gap-filling method can be used to readout cellular barcodes. Compared to the no-gap padlock approach, the gap padlock approach allows sequencing of the actual RNA sequence, but with lower sensitivity. The last approach, fluorescent *in situ* sequencing (10) (FISSEQ; Fig. 1C), can also be used to read out barcodes. In a recent version of FISSEQ (10), the cDNAs produced from reverse transcription of mRNAs are directly circularized using a single-stranded DNA ligase and used as templates for RCA. Because no prior knowledge of the mRNA is required, this direct-ligation approach can be used to sample the whole transcriptome *in situ*, but with even less sensitivity than the gap padlock approach. All three approaches utilize sequencing by ligation (9,10) to read out the sequences of the rolonies. Because only the gap padlock approach and the direct-ligation approach could be used for barcode sequencing *in situ*, both with low sensitivity, we sought to improve their sensitivity and optimize them for barcode sequencing.

**Figure 1.**
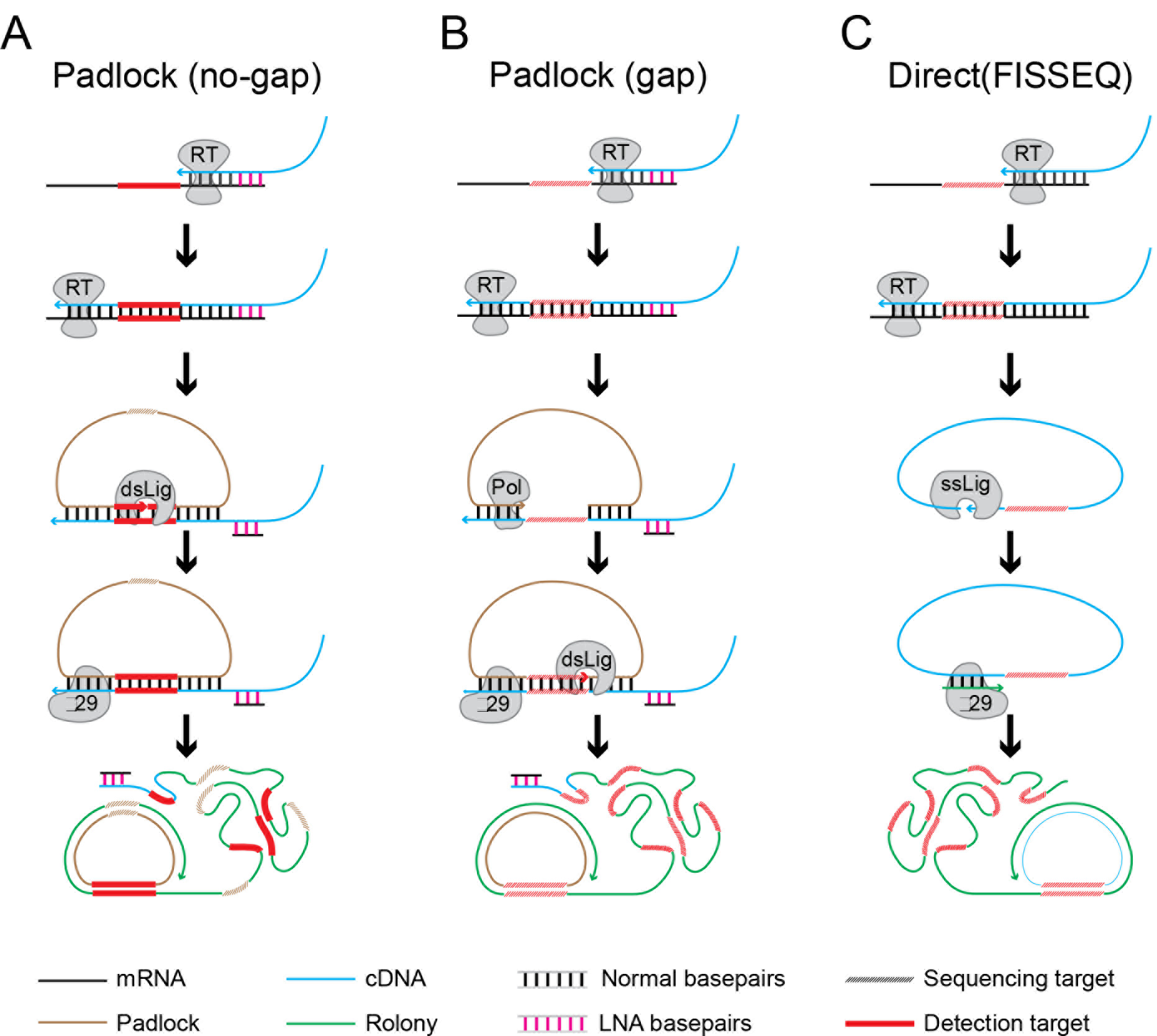
Comparison of *in situ* sequencing methods. In the padlock probe-based method without gap-filling (A), the RNA is reverse transcribed with an LNA primer (magenta) to produce a cDNA containing the sequence of interest (red). A padlock probe (yellow) containing a barcode (dashed) corresponding to the mRNA identity is hybridized onto the resulting cDNA and circularized with a double stranded DNA ligase. The circularized padlock probe is then used as a template for RCA and the barcode is sequenced *in situ*. In the padlock probe-based method with gap-filling (B), the two padlock arms hybridizes to the sequences flanking the area to be sequenced (red dashed). The padlock probe is then gap-filled with a DNA polymerase first, followed by circularization and RCA. In the direct-ligation based FISSEQ method (C), the RNA (black) containing the sequence of interest (red) is reverse transcribed, and the resulting cDNA (blue) is circularized with a single stranded DNA ligase. The circularized single stranded cDNA is then used as a template for rolling circle amplification (RCA) to generate a rolony (green).

Here we present BaristaSeq (<u>Bar</u>code *<u>i</u>n <u>s</u>itu* <u>ta</u>rgeted <u>seq</u>uencing), an improved version of the gap padlock probe-based method with a five-fold increase in efficiency, suitable for sequencing cellular barcodes similar to those used in MAPseq (6), a multiplexed barcode-assisted neuronal projection mapping method using next-generation sequencing. BaristaSeq is also compatible with Illumina sequencing by synthesis (SBS), which has a higher signal-to-noise ratio (SNR) than SOLiD sequencing. We demonstrate the accuracy and efficiency of BaristaSeq by sequencing random barcodes expressed in cells in culture.

## Material and Methods

### Viruses and oligonucleotides

All viruses and oligonucleotides used are listed in Supplementary Table S1 and S2. All oligonucleotides were acquired through IDT.

### Cell Culture

BHK cells were cultured in MEM-α (Thermo Fisher Scientific) supplemented with 5% FBS (Thermo Fisher Scientific), MEM Vitamin (Thermo Fisher Scientific), and Penn/Strep (Thermo Fisher Scientific) on glass bottom 96-well plates (Cellvis) coated with poly-D-lysine (Millipore). To infect BHK cells with Sindbis barcode libraries, 1μl diluted Sindbis virus (see Supplementary Table S1 for the dilutions for each virus) was added to a well in a 96-well plate containing freshly passed BHK cells. The cells were harvested 14-18 hours post-infection.

### *In vitro* gap-filling assay

*In vitro* gap-filling assays were performed in 1X Ampligase buffer (Epicentre) with 10 nM padlock probes (XC1149 and XC1151) and 10 nM cDNA template (XC1498), 20 μM dNTP, 0.012 U/μl Phusion High-Fidelity DNA Polymerase (Thermo Fisher Scientific) or Stoffel fragment (DNA Gdansk), additional 50 mM KCl, 20% formamide, and glycerol to a final concentration of 10%. The reaction was kept at 37 L for 30 min. We then terminated the reactions by adding 2x TBE Urea sample buffer (Thermo Fisher Scientific) and ran the samples on 15% Novex TBE-Urea gels (Thermo Fisher Scientific). The gel was then post-stained with SYBR-Gold (Thermo Fisher Scientific) in 0.5x TBE and imaged using a Canon xti dSLR camera.

For parameter optimization experiments, all conditions were kept the same as above except for the parameter being optimized.

To quantify band intensities, we de-noised the images using a median filter and subtracted the background with a 100 pixel radius rolling ball. We then quantified each band in ImageJ and two additional background areas in the same lane, one above all the bands and one below all the bands. We then subtracted the mean background intensities from all band intensities. Images shown in the figures were before the de-noising and background subtraction.

### BaristaSeq RNA amplification

Cells were washed in PBS first and then fixed in 10% buffered formalin (Electron Microscopy Sciences) for 30 mins followed by three washes in PBST (PBS with 0.5% Tween-20). The cells were then dehydrated in 70%, 85%, and 100% Ethanol, and were then kept in 100% Ethanol for at least an hour at 4°C. We then rehydrated the samples in PBST, treated the cells with 0.1M HCl for 5 mins at room temperature, followed by three washes in PBST. We then reverse transcribed the mRNAs overnight at 37°C with RevertAid H Minus M-MuLV Reverse Transcriptase (20 U/μl, Thermo Fisher Scientific) in 1X RT buffer supplemented with 1μM LNA RT primer, 500μM dNTP, 0.2μg/μl BSA and 1U/μl RiboLock RNase Inhibitor (Thermo Fisher Scientific).

On the second day, we washed the cells in PBST followed by crosslinking using 50mM BS(PEG)_9_ (Thermo Fisher Scientific) in PBST for 1 hour at room temperature. The crosslinkers were then washed out using 1M Tris-HCl, pH8.0, and the cells were further incubated in 1M Tris-HCl for 30 mins to neutralize residual crosslinkers. We then hybridized the padlock probes, gap-filled, and ligated them in an enzyme mix of 0.4 U/μl RNase H (Enzymatics), 0.2 U/μl Phusion High-Fidelity DNA Polymerase (Thermo Fisher Scientific), and 0.5 U/μl Ampligase (Epicentre) in 1X Ampligase buffer (Epicentre)
supplemented with 100 nM padlock probes, 50 μM dNTP, 1 U/μl RiboLock RNase Inhibitor, 50 mM KCl, and 20% formamide. The reactions were kept for 30 mins at 37°C followed by 45 mins at 45°C. We then washed the samples twice in PBST, and proceeded to rolling circle amplification at room temperature overnight using 1 U/μl ◻29 DNA polymerase (Thermo Fisher Scientific) in 1X ◻29 DNA polymerase buffer supplemented with 0.25 mM dNTP, 0.2 μg/μl BSA, 5% glycerol, and 20 μM aminoallyl-dUTP.

On the third day, we washed the cells in PBST, crosslinked the samples using BS(PEG)_9_ and neutralized the crosslinkers with 1M Tris-HCl. To visualize rolonies, we washed the samples twice with 2x SSC with 10% formamide, and incubated in the same buffer containing 0.5 μM fluorescent probes for 10 mins at room temperature, followed by three washes in 2x SSC with 10% formamide and one wash in PBST. To strip the fluorescent probes from the rolonies, we incubated the samples in 80% formamide that was preheated to 90°C for four times, each for two minutes. We then washed the samples three times in 2x SSC with 10% formamide. To hybridize sequencing primers, we washed the samples twice in 2x SSC with 10% formamide, followed by incubation in the same buffer with 2.5 μM sequencing primers for 10 mins at room temperature. The samples were then washed three times in 2x SSC with 10% formamide.

For *in situ* barcode amplification using the direct-ligation method, we followed the procedure described by Lee et al. (10) using a targeted RT primer (XC1016) and a targeted RCA primer (XC1017).

### BaristaSeq *in situ* sequencing

To sequence the samples using Illumina chemistry, we manually performed sequencing reactions using Illumina HiSeq SBS kit v4 (Illumina).

For the first cycle, we first washed the samples twice in SB2 at 60°C for three minutes, followed by two synthesis steps in IRM at 60°C for three minutes. We then washed the samples four times in PBST at 60°C, and then kept the sample in USM for imaging. After imaging, we briefly washed the samples twice in SB3, and then cleaved the fluorophores by incubating the samples twice in CRM for three mins at 60°C. We then briefly washed the samples twice in SB1 and twice in SB2, and proceed to IRM incubation as in the first cycle. When adapting this sequencing protocol to other samples, the timing required for each step likely varies with the physical format of the sample due to differences in the rate of heat transfer. The exact timing for each step should therefore be experimentally determined for a particular system to ensure that both the incorporation and the cleavage reactions are pushed to completion in each cycle.

To sequence the samples using SOLiD chemistry, we used the SOLiD FWD SR S50 Kit (Thermo Fisher Scientific) and followed the procedure described by Lee et al. (10). Briefly, we washed the samples twice with 1x Instrument Buffer (Thermo Fisher Scientific) and then incubated the samples in a reaction mix containing 6 U/μl T4 DNA ligase (Enzymatics) and 5 μl SOLiD sequencing oligonucleotides in 200 μl of 1x T4 DNA ligase buffer for 45 mins at room temperature. We then washed the samples four times with 1x Instrument Buffer for 5 mins each, and imaged the sample. After imaging, we incubate the samples twice in Cleave Solution 1 for 5 mins each, followed by two incubations in Cleave Solution 2.1 for 5 mins each. We then washed the samples three times with 1x Instrument Buffer and proceeded to the next cycle.

### Subsampling and quantification of rolonies

We mixed two padlocks (XC1613 and XC1614) at 1:100 ratio during the padlock gap-filling/ligation step. The two padlocks have the same arm sequences, but the backbone sequences differ so that they can be recognized by two different fluorescent probes (XC92 and XC1380). After rolony generation, we probed the samples with the probe targeting the diluted padlock and counted the rolonies. To prevent bias caused by the difference between the two padlock probes, we swapped the two padlock probes and counted the rolonies from the lesser padlock probe again. We then pooled the data and calculated the mean to get the 100-fold subsampled rolony counts.

The rolonies were counted in ImageJ using the “Find Maxima” function with a predetermined threshold across all samples. All p values reported were obtained using two-tailed Student’s t test with Bonferroni correction.

### Microscopy

All microscopy was performed on a Perkin-Elmer Ultraview Vox spinning disk confocal microscope or a Zeiss LSM 710 laser scanning confocal microscope. For sequencing images, the following settings were used for the four imaging channels on the LSM 710: G(cyan), 514nm Ex, 483-559nm Em; T(yellow), 514nm Ex, 635-693nm Em; A(magenta), 633nm Ex, 604-672nm Em; C(white), 633nm Ex, 700-758nm Em. The settings on the Ultraview Vox were: G(cyan), 514nm Ex, 550/49nm Em; T(yellow), 561nm Ex, 615/70 nm Em; A(magenta), 640nm Ex, 679/29nm Em; C(white), 640nm Ex, 775/140nm Em. The filter cubes on the Ultraview Vox were obtained through Semrock. The images shown in Fig. 4C and the subsampling images in Fig. 3A were acquired on the spinning disk microscope. All other images were obtained on the laser scanning confocal microscope.

To compensate for the bleed through of the four sequencing dyes in the four imaging channels and to normalize their signals, we generated rolonies using padlocks XC1308-XC1311 in four separate samples, and using a mixture of the four padlocks in a fifth sample, then sequenced one base using XC1312 as the sequencing primer. XC1312 reads into a barcode built into the backbones of the padlocks XC1308-XC1311, thus allowing the first four samples to contain sequencing signals from only one dye. Because the cells were filled with rolonies, the cells in the fifth sample contained a roughly equal amount of rolonies in all four dyes. We then imaged the first four samples using the four channels to calculate the bleed through, and used the fifth sample to adjust the exposure and gains to equalize the signals for the four sequencing dyes.

### Data analysis

All post-processing of sequencing data were done in Matlab (Mathworks). We first compensated for the bleed through in the four channel images. We then aligned the images from multiple sequencing cycles using Enhanced Cross Correlation(18) (ECC, available at http://iatool.net/). We manually selected ROIs on sequenced cells, further subtracted background caused by the nuclei, and assigned the channel with the strongest signal as the base in that cycle. The quality of the basecalls were defined as the intensity of the strongest channel divided by the root mean square of all four channels. Cells that expressed multiple barcodes were identified as cells with significant signals in two or more channels in more than two of the first five cycles.

The calculation of SNR was done in six fields of view in two biologically independent samples each for Illumina sequencing and SOLiD sequencing. In each field of view, we first merged the four sequencing channels into a single channel, taking the maximum value of the four channels at each pixel. We then calculated the SNR for three infected cells in that field of view by dividing the mean intensity in each cells against mean intensities of three uninfected cells, all corrected for the black point of the camera. The black point of the camera is measured as the mean value at three locations within the same field of view that had no cells or rolonies. The SNR of the field of view was taken as the mean of the SNR of the three cells, and the total SNR for that cycle was the mean SNR of the six fields of view.

### Results

In what follows we describe and characterize BaristaSeq. First, we demonstrate that aberrant gap-filling is an important mechanism limiting the sensitivity of previous *in situ* padlock-based sequencing approaches, and show that this inefficiency can be overcome by using a different polymerase. Next, we optimize BaristaSeq for sequencing long barcodes. Finally we characterize the sensitivity and accuracy of BaristaSeq by sequencing a 15-nt barcode in BHK cells in culture.

### Improved gap-filling in padlock-based targeted *in situ* RNA amplification

We first sought to identify the causes for the lower efficiency of the gap-filling padlock-based method. The padlock-based method can be used without gap-filling to detect mRNAs (Fig. 1B). The efficiency, however, drops significantly when gap-filling is used to interrogate the mRNA sequences. Because ligation of a gap padlock requires the polymerase to fill the gap exactly to the base next to the ligation junction (Fig. 2A, top), gap-filling could reduce the overall efficiency in two ways. First, insufficient polymerization would cause the gap to be filled only partially (Fig. 2A, middle), making ligation impossible. Second, some DNA polymerases with strand displacement activity may over-extend the padlock probe, also preventing the ligation (Fig. 2A, bottom).

**Figure 2.**
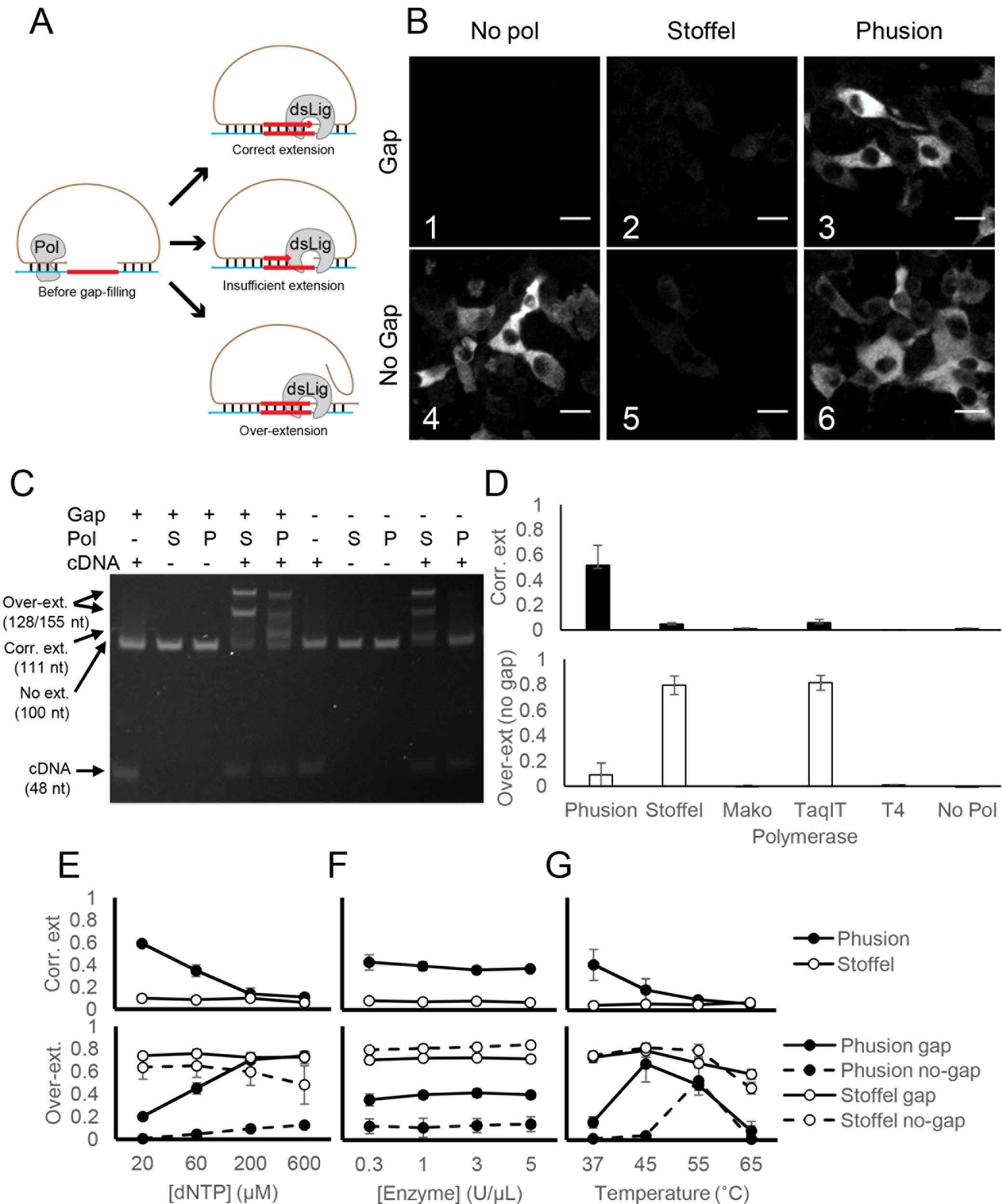
Optimization of gap-filling in *in situ* barcode sequencing. (A) Illustrations of normal (top) and aberrant gap-filling products caused by insufficient gap-filling (middle) and overextension (bottom). (B) *In situ* barcode amplification in infected BHK cells with padlock probes that require (Gap) or do not require (No Gap) gap-filling using the indicated polymerases. Scale bars = 20 μm. (C) *In vitro* gap-filling assay performed on padlocks that require (+ Gap) or do not require (− Gap) gap-filling using no polymerase (−), the Stoffel fragment (S), or Phusion DNA polymerase (P) in the presence (+ cDNA) or absence (− cDNA) of a cDNA template. Arrows on the left, from top to bottom, indicate positions for strand displaced gap-filling product, correct gap-filling product for “+ Gap” padlock, pre-gapfilling padlock, and the cDNA template. (D) The means and SEMs of the fraction of the correct gap-filling product using the gap padlock (top, black bars) and the fraction of over-extended products using the nogap padlock (bottom, white bars) with the indicated polymerases. N = 3 for all enzymes. (E)-(G) The means and SEMs of the faction of correct gap-filling product using the gap padlock (top) and the fraction of over-extended products (bottom) using either the gap padlock (solid lines) or the no-gap padlock (dashed lines) with either Phusion DNA polymerase (black dots) or the Stoffel fragment (white dots). The reactions were performed with the indicated dNTP concentrations (E), with the indicated enzyme concentrations (F), or at the indicated temperature (G). N = 3 for each condition in (E) and (F), and N = 4 for each condition in (G).

For *in vitro* padlock probe applications, many factors, including probe concentration, dNTP concentration (13), and polymerase choice(14), affect the total efficiency. Many DNA polymerases, including the Tfl polymerase (15), variants of the Taq polymerase (13,16,17), and variants of the Pfu polymerase (14), have been used for gap-filling. The Stoffel fragment polymerase, a truncated form of Taq polymerase without flap endonuclease activity, has been a popular choice for these applications and for *in situ* amplification of mRNA (9), presumably because of weaker strand displacement activity (17). It is unclear, however, whether the Stoffel fragment actually gap-fills effectively both *in situ* and *in vitro*.

To test the efficiency of gap-filling, we amplified a Sindbis virus RNA target using two different padlock probe designs, one with an 11-nt gap between the two padlock arms and one without a gap. As expected, in the absence of a gap-filling polymerase, the target RNA was efficiently amplified with a no-gap padlock, but no amplification was observed with the gap padlock (Fig. 2B1 vs Fig. 2B4). Consistent with previous reports, gap-filling using the Stoffel fragment polymerase on the gap padlock was inefficient (Fig. 2B2).

If the Stoffel fragment is inefficient at polymerization *in situ*, then we would not expect it to affect the overall efficiency when the no-gap padlock was used. Our results, however, did not support this hypothesis, because adding the Stoffel fragment polymerase when using the no-gap padlock—conditions under which no polymerization is required—actually reduced the efficiency of amplification (Fig. 2B5 vs Fig. 2B4) to a similar degree as when the gap padlock was used (Fig. 2B2). This suggested that the reduced efficiency observed when using the Stoffel fragment for gap-filling was not due solely to insufficient polymerization *in situ*.

We hypothesized that the Stoffel fragment was producing over-extended products. Such aberrant products could arise if the absence of flap endonuclease activity in the Stoffel fragment did not eliminate its strand displacement activity. We tested this hypothesis by performing the gap-filling reaction *in vitro* on a mock cDNA product. Consistent with the hypothesis that the padlock probes were overextended, most padlocks were extended either for a few bases or to the end of the cDNA when the Stoffel fragment was added to the no-gap padlock in the presence of the cDNA (Fig. 2C). No extension was observed in the absence of the cDNA, suggesting that such over-extension was template-dependent. Furthermore, the Stoffel fragment also overextended the gap padlock in the presence of cDNA without producing the correct gap-filling product (Fig. 2C). These results support the hypothesis that the strand displacement activity of the Stoffel fragment produced aberrant gap-filling products.

If strand displacement reduced the efficiency of gap-filling, then using a DNA polymerase without strand displacement activity should increase the efficiency of gap-filling. We therefore tested several DNA polymerases without strand displacement activity for gap-filling *in vitro* (Fig. 2D). Phusion DNA polymerase, a Pfu-based DNA polymerase, produced much less overextended product in the absence of a gap and converted 52 ± 2% (N = 13) of all gap padlocks to the correct gap-filled product (Fig. 2C). No other polymerase tested produced efficient gap-filling. We speculate that the inefficient polymerization seen with other polymerases might be due to the presence of formamide in our reactions, which is necessary for gap-filling *in situ*. We screened different dNTP concentrations (Fig. 2E), enzyme concentrations (Fig. 2F), and reaction temperatures (Fig. 2G) for the optimal reaction conditions for both Phusion polymerase and the Stoffel fragment. Compared to Phusion polymerase under the optimal conditions, the Stoffel fragment always underperformed in gap-filling efficiency (Fig. 2E-G, upper panel) and produced more over-extended products (Fig. 2E-G, lower panel) under all conditions we have tested. Consistent with the efficient gap-filling *in vitro*, Phusion DNA polymerase also resulted in efficient amplification of RNA *in situ* using both the gap padlock and the no-gap padlock (Fig. 2B). Therefore, Phusion DNA polymerase is more efficient at gap-filling on padlock probes than the Stoffel fragment due to the lack of strand displacement activity. We therefore used Phusion DNA polymerase for BaristaSeq.

To estimate the relative efficiency of BaristaSeq compared to existing methods, we generated rolonies on baby hamster kidney (BHK) cells infected with a Sindbis virus. Because the density of rolonies exceeded the optical resolution, we subsampled 1% of all rolonies (see Methods) and estimated the total number of rolonies. Briefly we used two padlock probes with the same sequences in the two arms complementary to the target sequence, but with different sequences in the backbone. The differences in the backbone allowed us to visualize each probe individually using one of two fluorescent oligonucleotides. To subsample the rolonies, we generated rolonies using a mixture of the two padlock probes at a ratio of 1:100, and visualized the rolonies with the fluorescent oligo complementary to the lesser padlock probe. We then estimated the total number of rolonies by multiplying the number of visualized rolonies by 100. The original padlock strategy generated 1300 ± 100 (mean ± SEM, n = 59; Fig. 3A, B) rolonies per cell, whereas our improved method generated 7000 ± 200 rolonies (n = 63, p < 0.0005 compared to the original strategy; Fig. 3A, B). In comparison, the targeted direct-circularization FISSEQ method (Fig. 1C) generated 12 ± 1 (n = 36, p < 0.0005 compared to both padlock strategies; Fig. 3A, B) rolonies. The false detection rate as measured in non-infected cells was also lower with the optimized padlock method [0.2 ± 0.1 rolonies per cell (n = 17)] compared to both the direct method [1.7 ± 0.4 per cell (n = 17), p < 0.005] and the original padlock probe-based approach (0.9 ± 0.2 rolonies per cell, n = 16, p = 0.01; Fig. 3C). We thus conclude that BaristaSeq was 5.4-fold more sensitive, and also more specific, than the original gap-filling padlock method using the Stoffel fragment.

**Figure 3.**
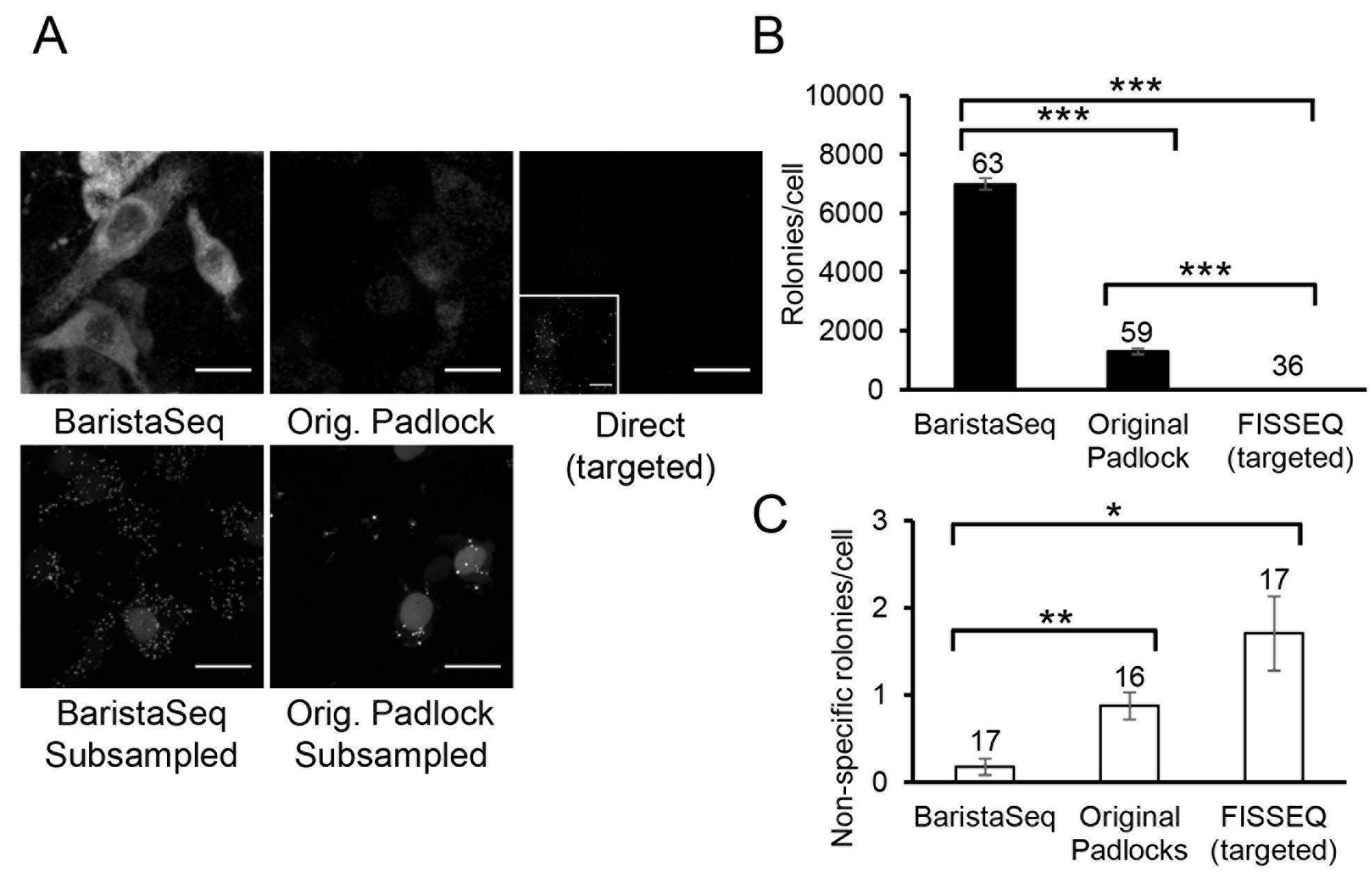
Quantification of rolony formation using BaristaSeq. (A) Top row: representative images of barcode amplicons in BHK cells using the indicated methods. Inset in the top right image shows the same image with 20x gain during post-processing. Bottom row: representative images of barcode amplicons in BHK cells with 100x subsampling. One in ~100 rolonies were visualized in these images. Scale bars = 20 μm. (B)(C) Average number of barcode amplicons per cell using the indicated methods in BHK cells with (B) or without (C) Sindbis virus. Error bars indicate SEM. *p = 0.01, **p < 0.005, ***p <0.0005 after Bonferroni correction. The numbers of cells counted are indicated on top of each bar. Quantification for both padlock approaches were done using subsampling.

### Adapting Illumina sequencing for *in situ* barcode sequencing

The original padlock approaches described by Ke et al.(9) were designed for reading SNPs and short barcodes (up to four bases) corresponding to gene identities. To sequence longer barcodes, we stabilized the rolonies through crosslinking using a method similar to that described previously (10). This modification stabilized the rolonies even under repeated heating and stripping with formamide (Fig. 4A, B).

**Figure 4.**
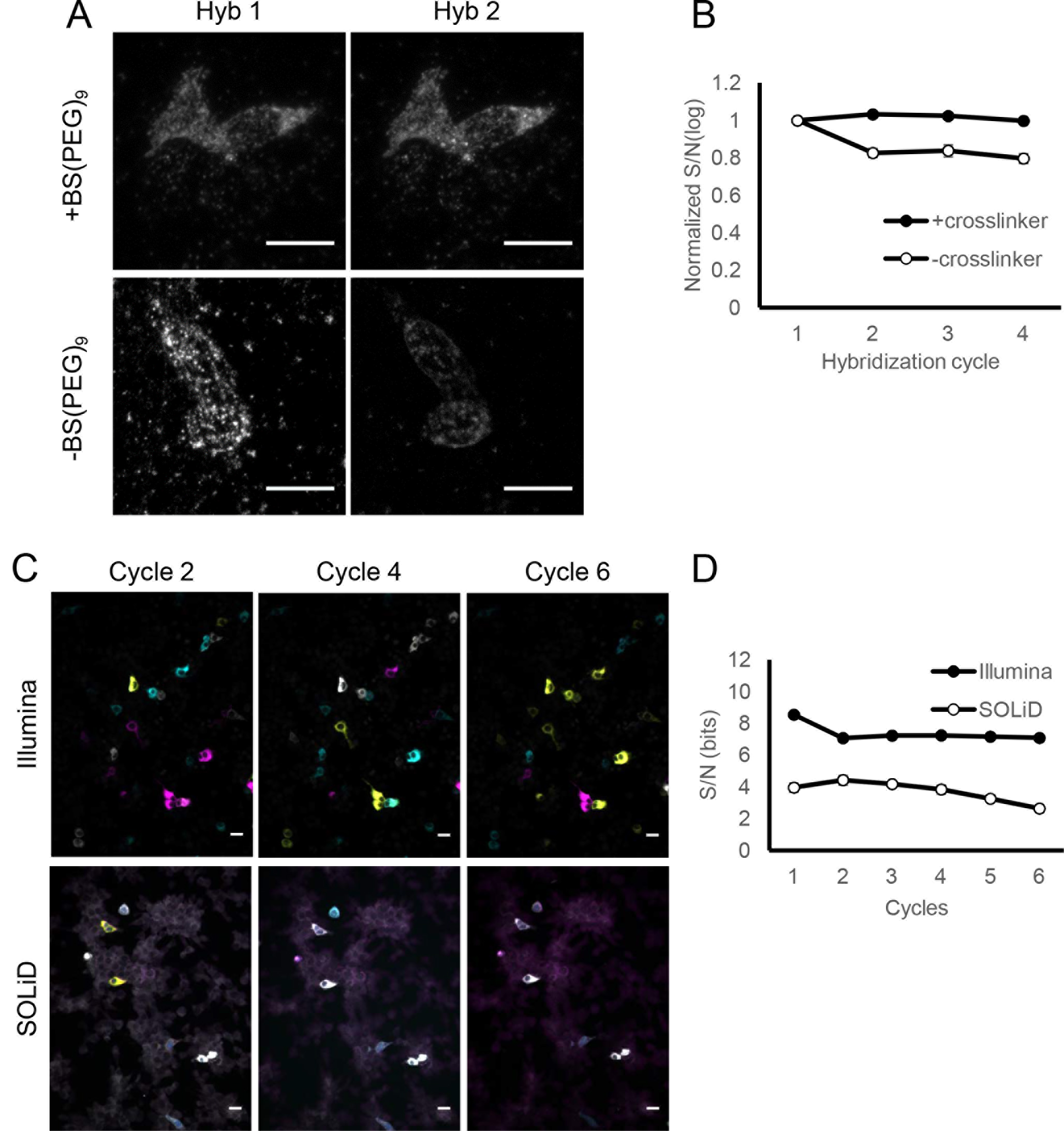
Improving *in situ* barcode amplification for sequencing. (A) Barcode amplicons with (+) or without (−) the post-RCA crosslinking before (Hyb 1) and after (Hyb 2) stripping with formamide and reprobing with a fluorescent probe. Scale bars = 20 μm. (B) Mean S/N ratios of rolonies during several hybridization cycles with (+) or without (−) BS(PEG)_9_ crosslinking after RCA. Error bars indicate SEM. N = 3 biological replicates. (C) Representative images of *in situ* barcode sequencing images using Illumina (top row) sequencing or SOLiD (bottom row) in BHK cells. The images were median filtered, but did not go through background subtraction. Scale bars = 20 μm. (D) Comparison of S/N ratio for SOLiD sequencing and Illumina sequencing over six cycles.

We also adapted Illumina sequencing chemistry to sequence rolonies generated using BaristaSeq. Previously reported *in situ* sequencing methods all utilized sequencing by ligation. We reasoned that due to multiple years of commercial optimization and the performance of Illumina sequencing chemistry *in vitro*, Illumina sequencing may produce better signals than sequencing by ligation. Indeed, sequencing by ligation using SOLiD chemistry produced stronger background fluorescence (Fig. 4C) and on average two-fold lower signal-to-noise ratio (Fig. 4D) than Illumina sequencing chemistry. Therefore, Illumina chemistry is superior to SOLiD for *in situ* barcode sequencing.

### Sequencing cellular barcodes in situ using Illumina sequencing chemistry

To evaluate the accuracy of basecalling cellular barcodes using BaristaSeq, we infected BHK cells with a Sindbis virus library encoding random 30-nt barcodes. The same virus library was used in Multiplexed Analysis of Projections by Sequencing (MAPseq)(6), a barcode assisted method that maps the single-cell projection patterns of thousands of neurons simultaneously. We then amplified barcodes and sequenced the first 15 bases of the barcodes *in situ* using Illumina sequencing (Fig. 5A, B). The sequencing quality was consistently high throughout all 15 cycles (Fig. 5C). The intensity of the signals was maintained with only minimal (13.9 ± 0.9 bits in cycle 1 vs. 11.9 ± 0.8 bits in cycle 15) reduction over 15 cycles (Fig. 5D). The fractions of the four bases remained approximately equal, as expected from a random barcode library (Fig. 5E). As expected, for each particular base call, the quality of the base call generally increased with the intensity of the signals (Fig. 5F).

**Figure 5.**
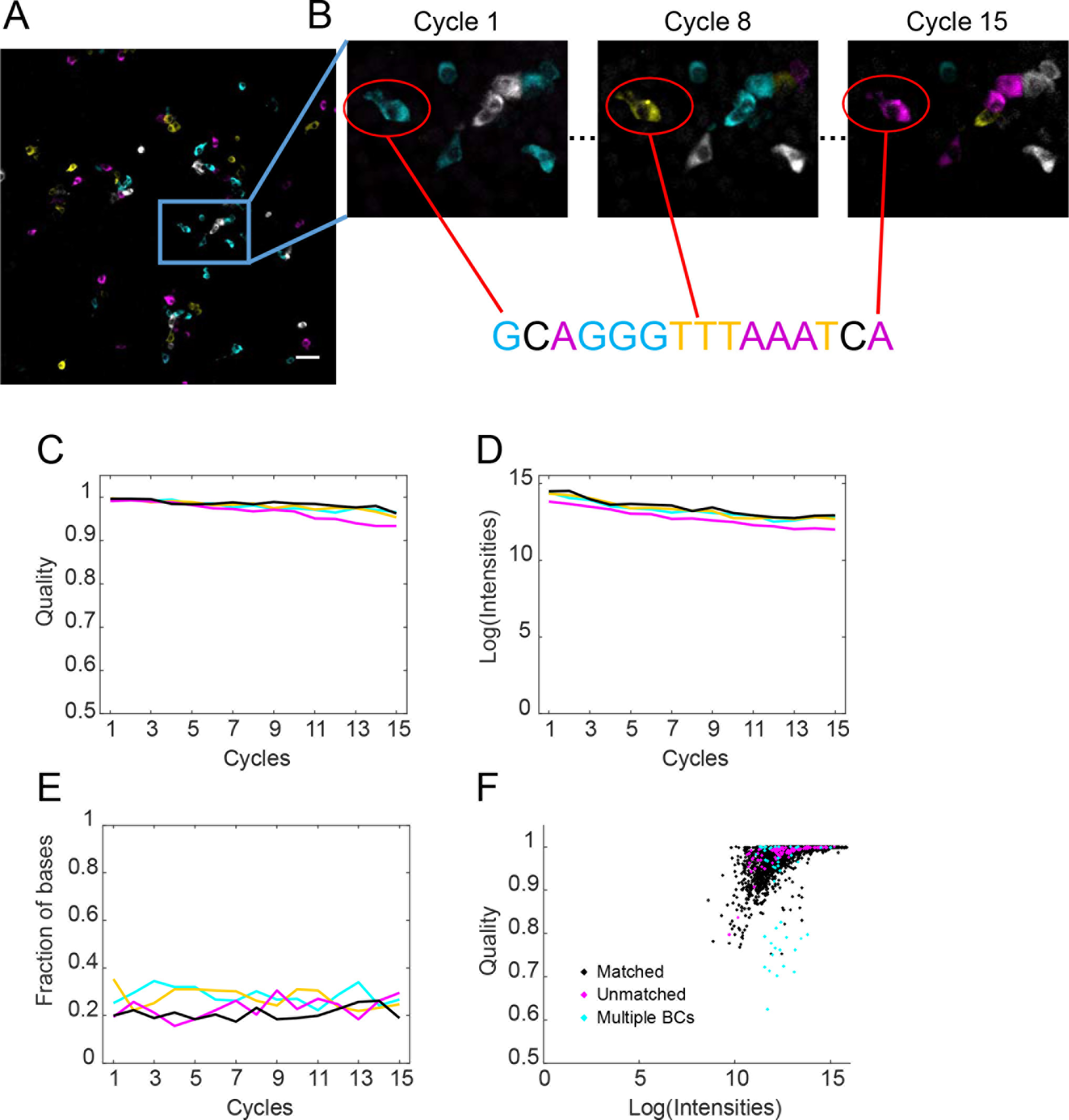
BaristaSeq in barcoded BHK cells. (A)Representative sequencing image of the first cycle. Scale bar = 50 μm. (B) Sequencing images of cycle 1, cycle 8, and cycle 15 of the indicated area from (A). The circled cell is basecalled below the images. (C) The intensity of the four channels, (D) fraction of the four bases, and (E) the quality of the base calls are shown over 15 cycles of Illumina sequencing *in situ*. (F) Quality of base calls and the intensity of the called channel for individual base calls. Black indicates correct base calls; magenta indicates base calls from *in situ* reads that did not match to *in vitro* reads; cyan indicates base calls from cells expressing more than one barcode.

We aligned *in situ* sequencing reads with all known barcode sequences in the library. The library contained 1.5 million known 30-nt barcode sequences, which represented ~97% of all barcodes in the library. For *in situ* sequencing, we called barcodes in 206 randomly selected cells from four imaging fields. Three cells were infected with more than one barcode per cell and were not analyzed further. Out of the remaining 203 cells, 97% (196) of called barcodes were perfect matches to sequences in the known barcode ensemble (Fig. 6A, B). To determine if these matched barcodes were due to chance, we generated random 15-nt barcodes and attempted to match them to the known barcode ensemble. Only 0.27 ± 0.05 (mean ± SEM, n = 100) out of 206 random 15-nt sequences matched perfectly by chance (typical random sequences had two mismatches), suggesting that the matched barcodes were unlikely to be false positives. The accuracy of BaristaSeq for sequencing 15-nt barcodes is thus at least 196/203, or 97%.

**Figure 6.**
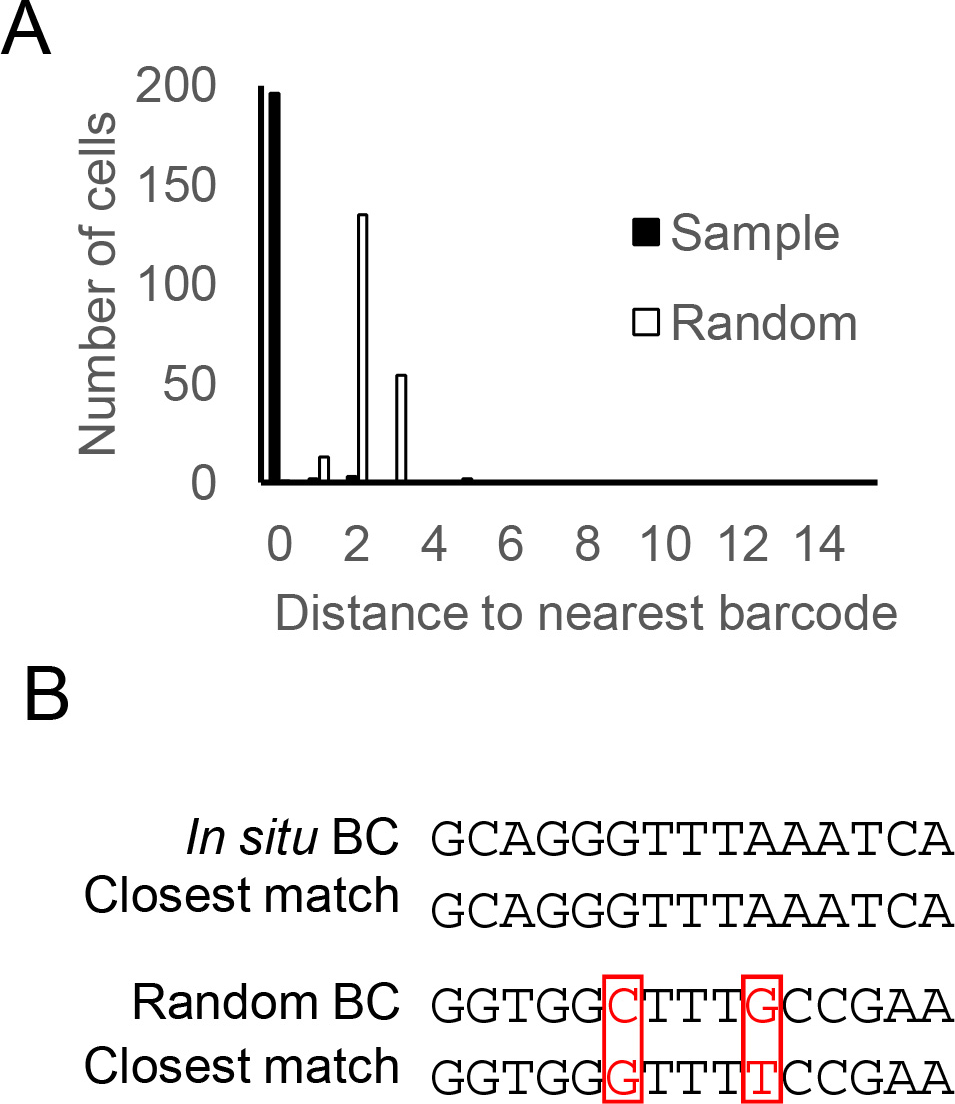
Comparison of BaristaSeq reads to conventional Illumina sequencing reads (A) Histogram of the number of mismatches between the *in situ* reads and their closest matches from *in vitro* reads (Sample) and the number of mismatches between random sequences and their closest matches from *in vitro* reads (Random). (B) Examples of a barcode read *in situ* and its closest match *in vitro*, and a random sequence and its closest match *in vitro*. Red indicates mismatches.

The seven barcodes that did not match to the known barcode ensemble could have originated from sequencing errors, or they could correspond to the 3% unknown barcodes in the library. Our results suggest that the latter explanation is more likely, because the quality of the unmatched barcode reads (magenta, Fig. 5F) were similar to that of the matched barcode reads (black, Fig. 5F). In contrast, cells that were labeled with more than one barcode (cyan, Fig. 5F) had significantly lower qualities for many base calls. The real accuracy of BaristaSeq is thus likely to be higher than the estimated 97%. These results indicate that BaristaSeq is highly accurate for cellular barcode sequencing.

## Discussion

We have presented BaristaSeq, a method for efficient and accurate sequencing of cellular barcodes *in situ*. We examined the cause of the low efficiency during padlock probe-based targeted *in situ* RNA amplification with gap-filling and replaced the Stoffel fragment with a Pfu based DNA polymerase. The lack of strand displacement activity of Pfu significantly increased the efficiency of gap-filling. The strand displacement activity, however, may not be the only cause for inefficient gap-filling, since gap-filling with Pfu still produced fewer rolonies than using a gapless padlock probe without a polymerase (Fig. 2B). One possible cause of this difference is that the Pfu polymerase has 3’→5’ exonuclease activity, which may digest the padlock probes and/or the cDNA product and thus reduce the amount of amplicons. Therefore, engineering of the gap-filling polymerase and optimization of the reaction conditions may further increase gap-filling efficiency.

The original padlock-based method was designed as a multiplexed *in situ* mRNA detection method (without gap-filling) or as a targeted SNP detection method (with gap-filling). In both cases, sequencing is limited to only a few bases (up to 4nt), and is performed at room temperature. The rolonies generated were stable enough under these conditions to allow sequence readout. To sequence longer barcodes, we made two modifications. First, we stabilized the amplicons through crosslinking (10). The increased stability of amplicons ensured minimal signal loss during heat cycles, which is essential for sequencing long barcodes. Second, we used Illumina sequencing chemistry instead of Sequencing by Ligation (SOLiD). We showed that Illumina sequencing has lower background signal than SOLiD when used *in situ*, even when sequencing short reads. One possible reason for this may be that the SOLiD reaction mix contains 1024 different types of short oligonucleotides, a quarter of which are labeled with a particular fluorophore, whereas the Illumina reaction mix only contains four different labeled nucleotides. The large number of unused oligonucleotides in each SOLiD cycle have the potential to bind non-specifically to tissue and increase background signal. Such differences may be more striking when sequencing in tissue sections, where non-specific binding of fluorophores tends to be more severe. Therefore, although both Illumina sequencing and SOLiD can sequence up to 35 nucleotides *in vitro* with similar performance, Illumina sequencing is superior for *in situ* sequencing.

DNA/RNA barcodes can be used as unique cellular tags to distinguish individual cells. DNA barcodes have been used in cellular lineage tracing for either small populations of cells (1–4) or throughout a whole organism (5). Recently, we have used RNA barcodes to label neurons for multiplexed projection mapping (MAPseq) (6). In these reports, the locations of the neurons under investigation had to be determined by manually picking individual neurons - a method which scales poorly. Efficient barcode sequencing *in situ* would allow us to preserve the location of each neuron’s soma while still maintaining high-throughput lineage tracing and projection mapping. Furthermore, in the case of MAPseq where the barcodes fill the neurons, *in situ* barcode sequencing would potentially allow visualization of the morphology of individual neurons. Preserving the locations of the cells being sequenced would allow correlation of lineages and projections with other information, such as gene expression assayed through FISH and neuronal activities assayed through functional imaging, at cellular resolution.

## Author contributions

X.C. and A.M.Z conceived the study. X.C. and Y.S. performed experiments. X.C. and A.M.Z. analyzed the data. X.C., A.M.Z., G.M.C., and J.H.L. discussed and wrote the paper.

## Data deposition

All *in situ* sequencing data are publicly available at https://www.dropbox.com/sh/f7p4yvy20tot5il/AAC6O5RnC7KlgcDC4YVxOgRZa?dl=0

## Acknowledgements

The authors would like to acknowledge Richie Kohman, Justus Kebschull, Ian Peikon, Ashlan Reid, and Shaina Lu for useful discussions and assistance, and Barry Burbach, Nour El-Amine, and Stephen Hearn for technical support.

## References

1. Walsh, C. and Cepko, C.L. (1992) Widespread dispersion of neuronal clones across functional regions of the cerebral cortex. Science, 255, 434–440.

2. Walsh, C. and Cepko, C.L. (1993) Clonal dispersion in proliferative layers of developing cerebral cortex. Nature, 362, 632–635.

3. Mayer, C., Jaglin, X.H., Cobbs, L.V., Bandler, R.C., Streicher, C., Cepko, C.L., Hippenmeyer, S. and Fishell, G. (2015) Clonally Related Forebrain Interneurons Disperse Broadly across Both Functional Areas and Structural Boundaries. Neuron, 87, 989–998.

4. Harwell, C.C., Fuentealba, L.C., Gonzalez-Cerrillo, A., Parker, P.R., Gertz, C.C., Mazzola, E., Garcia, M.T., Alvarez-Buylla, A., Cepko, C.L. and Kriegstein, A.R. (2015) Wide Dispersion and Diversity of Clonally Related Inhibitory Interneurons. Neuron, 87, 999–1007.

5. McKenna, A., Findlay, G.M., Gagnon, J.A., Horwitz, M.S., Schier, A.F. and Shendure, J. (2016) Whole-organism lineage tracing by combinatorial and cumulative genome editing. Science, 353, aaf7907.

6. Kebschull, J.M., Garcia da Silva, P., Reid, A.P., Peikon, I.D., Albeanu, D.F. and Zador, A.M. (2016) High-Throughput Mapping of Single-Neuron Projections by Sequencing of Barcoded RNA. Neuron, 91, 975–987.

7. Chen, K.H., Boettiger, A.N., Moffitt, J.R., Wang, S. and Zhuang, X. (2015) RNA imaging. Spatially resolved, highly multiplexed RNA profiling in single cells. Science, 348, aaa6090.

8. Moffitt, J.R., Hao, J., Bambah-Mukku, D., Lu, T., Dulac, C. and Zhuang, X. (2016) High-performance multiplexed fluorescence in situ hybridization in culture and tissue with matrix imprinting and clearing. Proc Natl Acad Sci U S A, 113, 14456–14461.

9. Ke, R., Mignardi, M., Pacureanu, A., Svedlund, J., Botling, J., Wahlby, C. and Nilsson, M. (2013) In situ sequencing for RNA analysis in preserved tissue and cells. Nat Methods, 10, 857–860.

10. Lee, J.H., Daugharthy, E.R., Scheiman, J., Kalhor, R., Yang, J.L., Ferrante, T.C., Terry, R., Jeanty, S.S., Li, C., Amamoto, R. et al. (2014) Highly multiplexed subcellular RNA sequencing in situ. Science, 343, 1360–1363.

11. Larsson, C., Koch, J., Nygren, A., Janssen, G., Raap, A.K., Landegren, U. and Nilsson, M. (2004) In situ genotyping individual DNA molecules by target-primed rolling-circle amplification of padlock probes. Nat Methods, 1, 227–232.

12. Larsson, C., Grundberg, I., Soderberg, O. and Nilsson, M. (2010) In situ detection and genotyping of individual mRNA molecules. Nat Methods, 7, 395–397.

13. Li, J.B., Gao, Y., Aach, J., Zhang, K., Kryukov, G.V., Xie, B., Ahlford, A., Yoon, J.K., Rosenbaum, A.M., Zaranek, A.W. et al. (2009) Multiplex padlock targeted sequencing reveals human hypermutable CpG variations. Genome Res, 19, 1606–1615.

14. Shen, P., Wang, W., Chi, A.K., Fan, Y., Davis, R.W. and Scharfe, C. (2013) Multiplex target capture with double-stranded DNA probes. Genome Med, 5, 50.

15. Abravaya, K., Carrino, J.J., Muldoon, S. and Lee, H.H. (1995) Detection of point mutations with a modified ligase chain reaction (Gap-LCR). Nucleic Acids Res, 23, 675–682.

16. Porreca, G.J., Zhang, K., Li, J.B., Xie, B., Austin, D., Vassallo, S.L., LeProust, E.M., Peck, B.J., Emig, C.J., Dahl, F. et al. (2007) Multiplex amplification of large sets of human exons. Nat Methods, 4, 931–936.

17. Akhras, M.S., Unemo, M., Thiyagarajan, S., Nyren, P., Davis, R.W., Fire, A.Z. and Pourmand, N. (2007) Connector inversion probe technology: a powerful one-primer multiplex DNA amplification system for numerous scientific applications. PLoS One, 2, e915.

18. Evangelidis, G.D. and Psarakis, E.Z. (2008) Parametric image alignment using enhanced correlation coefficient maximization. IEEE Trans Pattern Anal Mach Intell, 30, 1858–1865.

